# Intraspecific genome size variation in *Rorippa indica* reveals a tropical adaptation by genomic enlargement

**DOI:** 10.1101/2024.10.13.618028

**Authors:** Ting-Shen Han, Quan-Jing Zheng, Jun-Xian Lv, Sisi Li, Zhi-Qiang Du, Yao-Wu Xing

## Abstract

Genome size exhibits substantial variation across organisms, but the underlying causes and ecological consequences remain unclear. While interspecific comparisons have suggested selective pressures against large genomes, intraspecific variation has been less explored. Here, we investigate genome size variation within the hexaploid yellowcress herb, *Rorippa indica*, by integrating flow cytometry, plastomic phylogeography, genomic repeat profiling, and reciprocal common garden experiments. We analyzed 192 accessions from 83 natural populations, revealing a 128 Mb range in genome size. Plastome haplotype analysis identified a haplogroup that shifted niches to tropics and evolved larger genomes. A strong correlation was found between larger genomes and tropical habitats characterized by higher temperatures and lower seasonality. Genomic repeat content, particularly 45S rDNA and *Ty1*-*copia* transposable elements, was associated with larger genomes. Reciprocal transplantation experiments confirmed the adaptive nature of large genomes in tropical environments, with individuals exhibiting lower growth rates but higher fecundity. Our findings support the “large genome selection” hypothesis, suggesting that genome size enlargement, driven by genomic repetitive elements, can be an adaptive response to high temperatures of tropics. As global warming continues, plants with larger genomes may exhibit slower growth but increased reproductive output, potentially impacting ecosystem dynamics and agricultural productivity.

## Introduction

A well-known phenomenon about biodiversity is that genomes come in surprisingly diverse packages across organisms. From the microscopic bacterium *Nasuia deltocephalinicola* with a mere 112,000 base pairs to the *Paris japonica* plant boasting 150 billion base pairs, genome size exhibits a remarkable range (Hidalgo *et al*., 2017). Such natural variation plays a crucial role in various biological processes, influencing genomic plasticity (Lynch & Conery, 2003; Leitch & Leitch, 2008), cellular dynamics (Beaulieu *et al*., 2008; Francis *et al*., 2008), environmental adaptation (Knight & Ackerly, 2002; Faizullah *et al*., 2021), and species diversification (Simonin & Roddy, 2018; Bhadra *et al*., 2023; Gomez *et al*., 2024). Although empirical evidence is scarce, it seems that adjustable genome size would be a feasible way to balance between relatively low mutation rate and demands of prompt response to changing environments (Mei *et al*., 2018). However, the dynamics of genome size variation remain largely shrouded in ambiguity, especially for its evolutionary trajectory and fitness effects under natural context.

The variation in genome size has significant evolutionary and ecological implications (Bennett, 1987), whereas the debate persists on whether genome size variation is driven by neutral evolution or natural selection. Two main neutral hypotheses—mutational hazard and mutational equilibrium—suggest genome size is shaped by random drift rather than selection (Petrov, 2002; Lynch & Conery, 2003). In contrast, adaptive hypotheses argue that genome size affects phenotypic traits such as cell size and metabolism (Beaulieu *et al*., 2008), which provides fitness effects under specific environments (Mei *et al*., 2018; Faizullah *et al*., 2021). Recently, accumulated evidence uncovered the adaptive aspects of genome size variation. Among them, the natural variation in genome size has been attributed to multiple evolutionary or ecological factors. Besides whole-genome duplication or polyploidy being a unique driver, intrinsic factors like dynamics of repetitive or transposable elements were largely explored (Bourque *et al*., 2018), as well as other ones, including environment (Kang *et al*., 2015; Bureš *et al*., 2024), mating system (Roessler *et al*., 2019), or demographic history (Lefebure *et al*., 2017; Bilinski *et al*., 2018). These results collectively suggest that genome size may function as an ecologically important trait to be targeted by natural selection.

Previous studies, primarily focused on interspecific comparisons, have yielded several hypotheses to explain the adaptive expression of genome size variation (Bureš *et al*., 2024). For example, the “large genome constraint” hypothesis suggests an evolutionary constraint for larger genomes and selective advantage of genome size shrinkage (Knight, 2005; Hidalgo *et al*., 2017; Heckel, 2020; Gomez *et al*., 2024). In addition, the “environmentally selected” hypothesis proposes a crucial role for genome streamlining in response to stresses, such as drought, intertidal changes, or limited nutrient availability (Lyu *et al*., 2018; Greenhalgh *et al*., 2020; Faizullah *et al*., 2021). Intriguingly, genomic enlargement was also observed to participate in evolutionary events, such as recent demographic fluctuation (Lefebure *et al*., 2017) or parallel altitudinal adaptation (Díez *et al*., 2013). For example, studies show that larger genomes are convergently found in stable environments with lower temperature seasonality and longer growing seasons (Knight & Ackerly, 2002; Díez *et al*., 2013; Carta & Peruzzi, 2016; Qiu *et al*., 2019). This may result from the relaxed constraint on growth rates and/or temperature-spurred activity of transposable elements, thus allowing larger-genome plants to thrive under warmer environments (hereafter, termed as the “large genome selection” hypothesis). Therefore, it is no doubt that either shrinkage or enlargement in genome size would be envisioned as the proxy of natural variation. However, their potential ecological and evolutionary significance are still awaiting to be fully untangled (Mei *et al*., 2018; Blommaert, 2020), particularly by attaining more insights from intraspecific investigations on larger genomes. Such studies at the population level could clarify the main forces driving natural variation in genome size.

This study explores the intriguing realm of intraspecific genome size variation. We shift focus from interspecific comparisons to uncover the fascinating phenomenon among populations of *Rorippa indica*, an edible and ecologically diverse species (Gu & Hsu, 1986; Tu *et al*., 2019; Castillo-Lorenzo *et al*., 2024). Plants of *R. indica* are natively distributed in temperate East Asia, and recently colonized into tropical areas, such as Southeast Asia or Central America (Setyawati *et al*., 2015). The species has an allohexaploid origin (2*n*=6*x*=48, AACCBB) between tetraploid *Rorippa dubia* (2*n*=4*x*=32, AACC) and diploid *Rorippa globosa* (2*n*=2*x*=16, BB) (Han *et al*., 2024), thereby allowing for the observation of pronounced genome size variation due to potential genomic redundancy. Its broad habitat range and phenotypic diversity also offers a unique opportunity to explore the ecological and evolutionary consequences of intraspecific genome size variation. Here, we aim to investigate the “large genome selection” hypothesis and address: (1) what is the extent and pattern of genome size expansion (if any) in natural populations of *R. indica* and what factors influence this variation; (2) does larger genome in *R. indica* correlate with fitness-related traits; and (3) what is the evolutionary trajectory of genome size in *R. indica*. By unraveling these questions, this study will shed light on drivers and consequences of genome size variation across natural populations. This knowledge will significantly contribute to our understanding of evolutionary mechanisms and how organisms adapt to their environments under global climate changes.

## Material and method

### Population sampling

A total of 192 accessions from 83 natural populations were sampled during 2017 to 2019 (Figure 1a), excluding the closely related species *R. hengduanshanensis* (Zheng *et al*., 2021). In 2020, 8‒10 seeds for each accession were stratified on ½ MS medium under 4℃ and dark for two weeks, and then germinated under 22℃ and long day conditions (16h light + 8h dark) for ten days. Seedlings with at least two true leaves were transplanted into mediums mixed with nutrient soil and vermiculite (1:3) under 22℃ and long day condition (16h light + 8h dark). To reduce systematic and artificial influences on the following cytological experiment, all plants were randomized into trays of 4 ×8 grids using the default method in R package agricolae v.1.3-3.

**Fig. 1.**
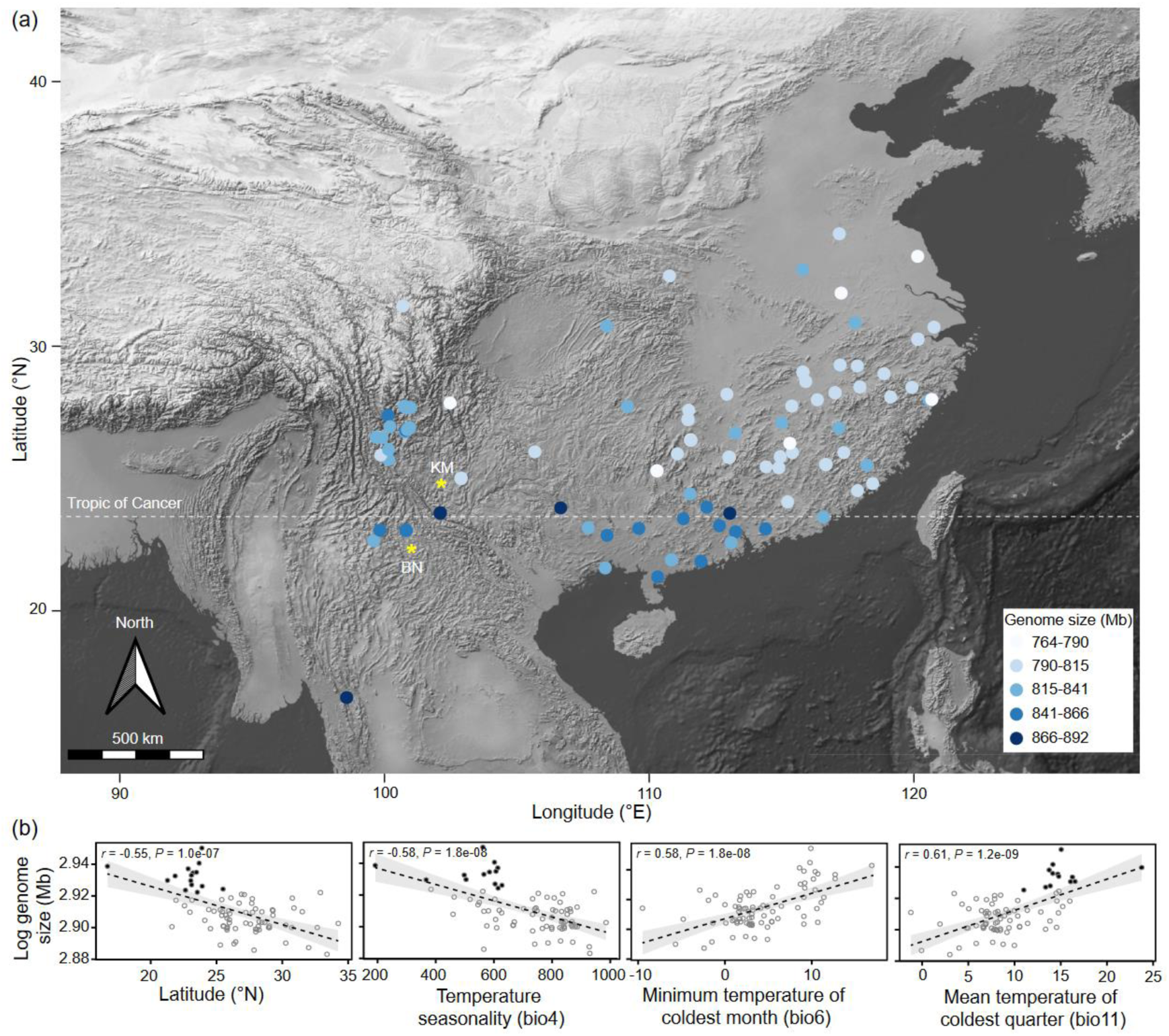
Natural variation in genome size of *R. indica*. (a) Geographic distribution of genome sizes in *R. indica*. The genome size is represented by the color of each accession dot, ranging from light (smaller genome) to dark blue (larger genome). Yellow asteroids indicate the locations of two common gardens: Kunming (KM) and Xishuangbanna (BN). (b) The relationship between log- transformed genome size and several environmental factors: latitude, temperature seasonality (bio4), minimum temperature of coldest month (bio6), and mean temperature of coldest quarter (bio11). In each panel, the dashed line represents the best-fitting linear regression model; the grey zone indicates the 95% confidence intervals for the regression line; and the Pearson’s correlation coefficient (*r*) and *P*-values are shown above.

### Genome size estimation

Flow cytometry (FCM) was used to measure genome size variation in *R. indica*. Fully developed fresh leaves were prepared according to the published method (Han *et al*., 2022), with plants of *Capsella rubella* MTE (219 Mb) as the internal standard. To accurately distinguish the G1 and G2 peaks of *R. indica*, FCM tests were performed with or without *C. rubella* leaves for each sample, separately. A total of 3‒ 4 replicates per accession were sampled and tested at two or three different developmental stages. Genome size was estimated for mixed samples as the total nuclear DNA content (i.e., 2C value) according to formula: sample genome size = (sample mean value of G1 peak / internal standard mean value of G1 peak) ×219 Mb. Estimates from multiple outputs of FCM experiments were extracted and combined using custom R script. The FCM experiment was performed using a BD FACSVerse Flow Cytometer (USA). To check the chromosome numbers of *R. indica* plants, slides were prepared for selected root samples representing different genome sizes (Han *et al*., 2015).

### Correlation with climatic variables

Bioclimatic variables (bio1‒19) for the 83 natural populations were extracted from WorldClim (https://www.worldclim.org/) using the R package Raster v.2.6-6. Log-transformed median values of intrapopulation genome size were used to estimate the Pearson’s correlation (*r*) with each bioclimatic or geographic variable (i.e., latitude, longitude, and elevation) in R v.4.0.2.

### Genomic sequencing and chloroplast genome assembly

High-quality DNA was extracted for 64 randomly selected accessions spanning the native range of *R. indica*. Raw whole-genome sequencing was performed on the Illumina Novaseq platform. Chloroplast genome assembly and annotation were performed by GetOrganelle v.1.4.0 (Jin *et al*., 2020) and CPGAVAS v.2.0 (Shi *et al*., 2019), respectively. Protein coding genes (PCGs) were extracted and concatenated using custom R script (Han *et al*., 2024). Sequences were aligned using MAFFT method in Geneious Prime v.2019.0.3 (Kearse *et al*., 2012).

### Phylogeny and network analysis

Chloroplast haplotypes were inferred from the aligned PCGs sequences using DnaSP v.6.10.04 (Rozas *et al*., 2017). Haplotype networks were built under TCS method using PopART v.1.7 (Leigh & Bryant, 2015), with gaps as the fifth variant and the limit of statistical parsimony connection as 95%. Haplotypic phylogeny was constructed using BEAST v.2.6.0, under relaxed molecular clock and uniform prior distributions (Han *et al*., 2024). A total of four independent runs were carried out, each with 200 million Markov Chain Monte Carlo simulations. Convergence was ensured by the effective sample size of parameters larger than 200 using TRACER v.1.7. Tree files were combined in LogCombiner v.2.6.0 (BEAST Developers), with 10% of runs as burn-in. Based on a set of 1000 resampled trees, a final maximum clade credibility tree was annotated under mean node heights in TreeAnnotator v.1.8.4 (BEAST Developers). The resampled trees were dated using penalized and maximum likelihood methods in R package ape v.5.5 (Paradis & Schliep, 2019), under the selected relaxed clock model. The maximum and minimum crown ages of *R. indica* were assigned as 1.474 and 0.712 Myr, respectively, according to the published dating results using a plastome tree (Han *et al*., 2024). The mean node ages were calculated in ape v.5.5, as well as their corresponding values of 95% highest posterior density (HPD). Integrated evidence for closely related haplotypes was clarified as haplogroup (HG) for subsequent analysis. Specifically, the assigned haplogroups may have been separated from each other by accumulated plastid variation, which were confirmed by both robust phylogenetic supports (i.e., posterior probabilities = 1.0) and convergent network clusters among haplogroups than those within haplogroup.

### Phylogeny-corrected correlation with climatic variables or repeat elements

To test the effect of climatic variables or repeat elements on genome size variation without phylogenetic inference, multiple phylogenetic generalized least squares (PGLS) tests were implemented using the R package Caper v.1.0.1 (Orme *et al*., 2013). Climatic variables (bio1‒19) were obtained as described above. To characterize the repetitive DNA from unassembled sequences, pair-ends raw reads under genomic coverage of 0.1×were sampled and merged from Illumina sequencing data for each sample using seqtk (https://github.com/lh3/seqtk). Genomic repeats were captured using the protein domain database of Viridiplantae v.3.0 in RepeatExplorer2 (Novak *et al*., 2020).

### Common garden experiment

To evaluate fitness effect of genome size (GS) variation, reciprocal transplantation experiment was performed for accessions with small (< 840 Mb) or large (> 840 Mb) genome size. Types of genome size were divided according to comparison with that of the constructed ancestral haplotype (H01 or HG-Ⅰ, see the result section for details). Two common gardens were established in temperate Kunming (KM; mean temperature of winter = 5‒10℃) and tropical Xishuangbanna (BN; 15‒20℃), respectively (Figure 1a). To reduce the influence of phylogenetic interference, accessions from one specific haplogroup (HG-Ⅲ) with varying genome sizes were used (Figure S4), including 11 accessions with small genome sizes and nine with large ones. The experimented seeds were collected from plants used for FCM and growing under the chamber for one generation. They were randomized into trays of 4 ×8 grids using the default method in R package agricolae v.1.3-3, with 15 biological replicates per accession per site. About 10 seeds per replicate were sowed directly into each grid with a medium made of nutrient soil to vermiculite = 1:3. Only one seedling was kept after germination per grid. The experiments were performed from October 2021 to August 2022, to cover the natural growing periods of *R. indica* during the autumn-spring seasons in temperate areas and the dry season in tropical areas.

A total of 22 traits were collected across the whole life history of *R. indica* plants. The original traits included the germination rate recorded at seven days after sowing (GR), survival rates at 10/14/18 weeks (SR_10W/14W/18W), mean or maximum rosette diameters at 10/14/18 weeks (RD_10W/14W/18W_mean/max), flowering time (FT), total number and total/mean length of primary inflorescence (TN/TL/ML_PI), the tallest length of primary inflorescence as plant height (PH), total number and total/mean/maximum length of secondary inflorescence (TN/TL/ML/MaxL_SI), and total number of fruits (TN_F). The calculated traits included the mean or maximum relative growth rates (RGR_mean/max) evaluated as the averaged increasement of rosette diameters per four-week. The effects of site (KM vs. BN), genome size/GS type (small vs. large), or their interaction (site ×GS) on trait variation were tested using generalized linear mixed model (GLMM) in the R package glmmTMB v.1.9.4 (Brooks *et al*., 2017). The influence of random effects (population and tray) was determined by our published method (Du *et al*., 2024).

## Results

### Genome size variation in *Rorippa indica*

According to FCM tests on 192 accessions from 83 natural populations, a large intraspecific variation in genome size was found in *R. indica* (Figure 1a). The mean (± standard deviation/SD) genome size for *R. indica* plants was estimated as 811.707 ± 36.741 Mb, ranging from 764 to 892 Mb. This is the largest variational range of genome size observed among *Rorippa* species (Figure S1), compared with the published SD values of 11.219 Mb in *R. elata* (186 accessions) or 10.779 Mb in *R. palustris* (62 accessions) (Han *et al*., 2022). Chromosome numbers were counted as 2*n*=6*x*=48 for the tested samples of *R. indica* with different genome sizes. Correlation analysis unveiled compelling connections between genome size variation and multiple geographic or bioclimatic variables (Figure S2). The top four environmental variables genome sizes correlated with were latitudes (*r* = −0.55, *P* = 1.0e-07), temperature seasonality (bio4, *r* = −0.58, *P* = 1.8e-08), the minimum (bio6, *r* = 0.58, *P* = 1.8e-08) and mean temperatures of winter (bio11, *r* = 0.61, *P* = 1.2e-09) (Figure 1b). In summary, plants of *R. indica* exhibited a high level of climate-related natural variation in genome size.

### Phylogeography of genome size variation

A total of 28 chloroplast haplotypes were inferred out of the sequenced 64 accessions. Based on the haplotypic tree and network, these haplotypes can be clustered into five haplogroups and coded as HG Ⅰ-Ⅴ (Figure 2a & S3).

**Fig. 2.**
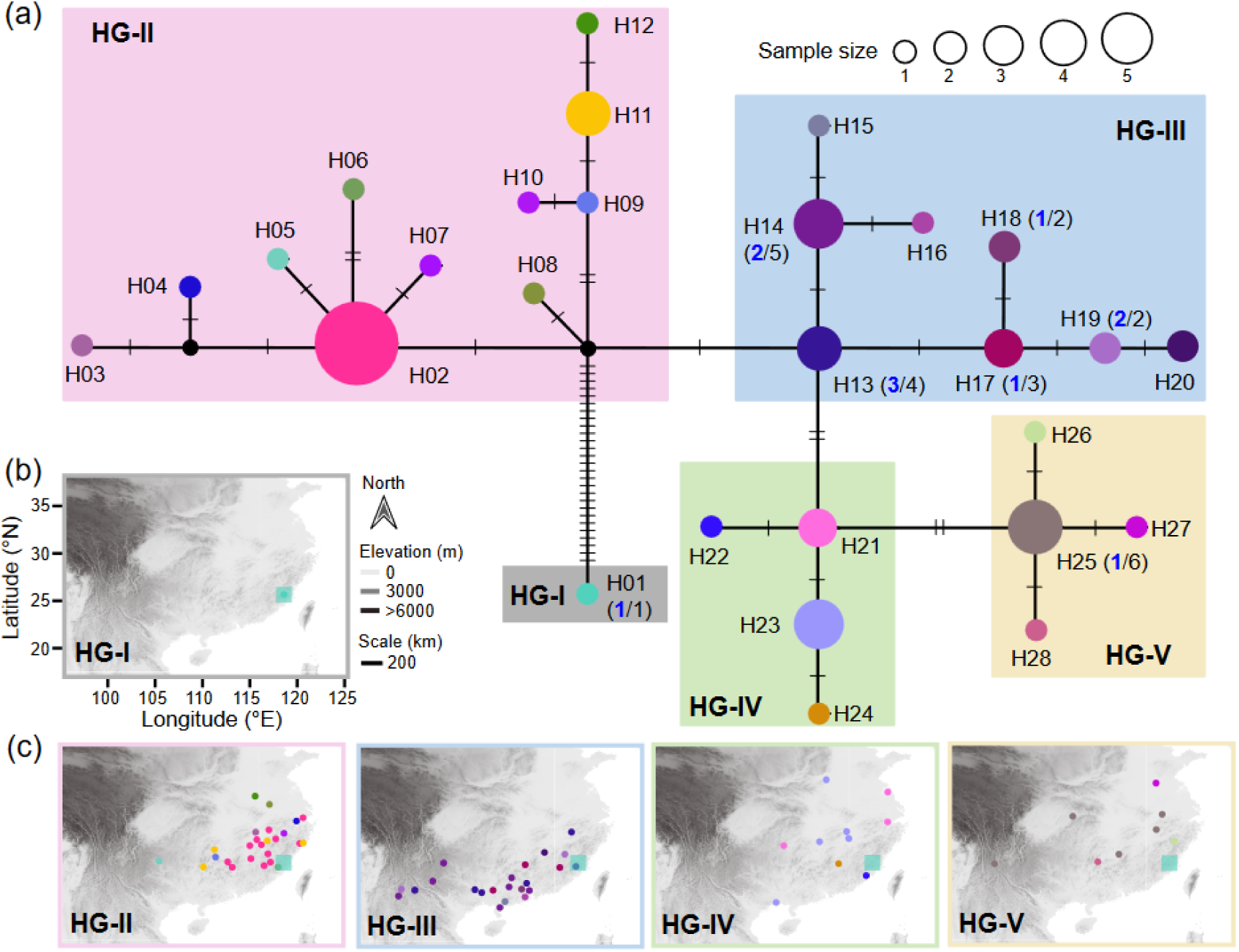
Phylogeography of plastome haplotypic diversity in *R. indica*. (a) Network of plastid haplotypes in *R. indicia*. Haplotypes are represented as colored pie charts, with names (H01‒28) and relationships indicated by connecting lines. The number of mutations between haplotypes is shown by bars on connecting lines. The size of each pie chart corresponds to the number of individuals with that haplotype. Haplotypes with larger genomes are labeled with blue numbers out of the total haplotype size indicated in parentheses. Haplogroups are labeled as HG Ⅰ-Ⅴ under shades with different colors. (b) Map of haplotypes from the ancestral HG-Ⅰ. (c) Maps of haplotypes from the derived HG-Ⅱ, HG-Ⅲ, HG-Ⅳ and HG-Ⅴ, respectively. In (b‒ c), haplotypes are illustrated as colored dots same with (a), with the position of HG-Ⅰ marked as a square in each panel.

Chloroplast haplogroups of *R. indica* harbored different geographic regions and genome sizes. For example, HG-Ⅰ has haplotypes (H01) sampled from southeast China (25.497°N, 118.231°E) (Figure 2b). HG-Ⅰ was rooted in the haplotypic tree and originated *c.* 1.400 million years ago (Ma) (95% HPD: 1.474‒1.144 Ma), differentiating from all the other samples and representing an ancestral haplogroup along phylogeny of *R. indica* plants (Figure 2a). The rest haplogroups HG Ⅱ-Ⅴ were derived from HG-Ⅰ and clearly diverged from each other. Compared with the geographic location of HG-Ⅰ, plants of HG-Ⅲ were significantly distributed southward into lower latitudes (mean ±SD = 24.051 ±2.069 °N; one-sample Wilcoxon test, *V* = 29, *P* = 0.012) since *c.* 1.071 Ma (95% HPD: 1.155‒0.902 Ma) (Figure 2c & 3a). In contrast, most plants of the rest three haplogroups (HG-Ⅱ, −Ⅳ, and −Ⅴ) clearly inhabited northward into higher latitudes than that of HG-Ⅰ (27.885 ± 2.069 °N; *V* = 899, *P* = 2.490e-06) (Figure 2c & 3a), after *c.* 1.164 (1.247‒0.974), 0.857 (0.956‒0.751), and 0.930 (1.000‒0.783) Ma, respectively.

**Fig. 3.**
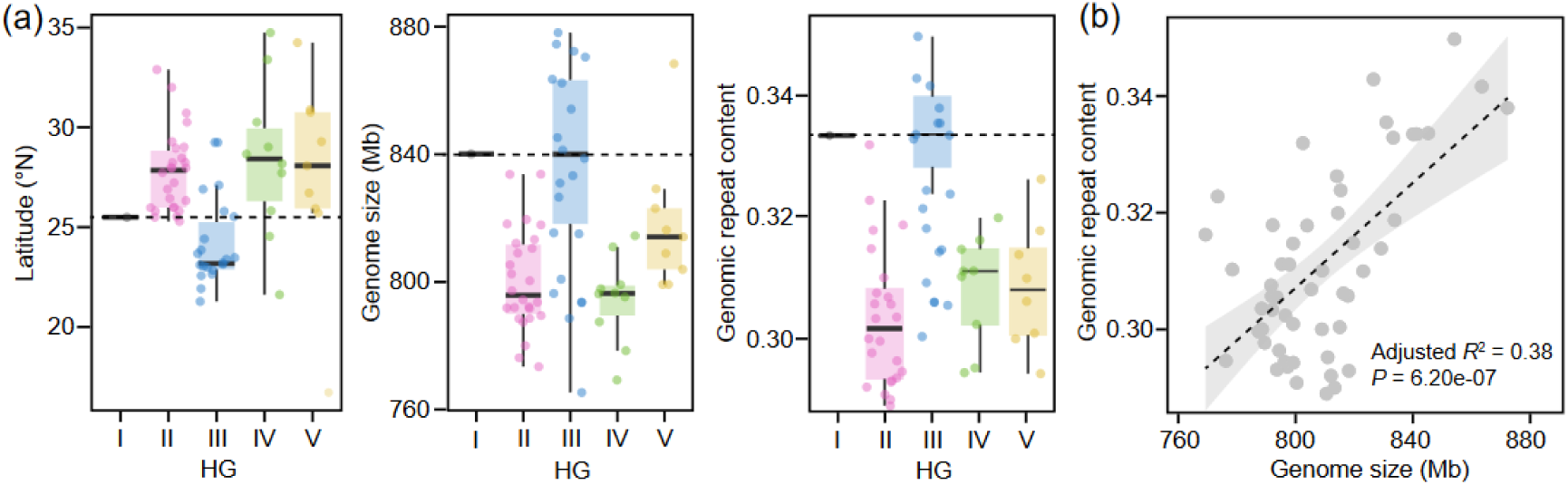
Geographic and genomic components of genome size variation. (a) Box-and-whisker plots illustrate the distribution of latitude, genome size, and genomic repeat content for each haplogroup (HG Ⅰ-Ⅴ). The median value is represented by the horizontal line within each box. The 25^th^ and 75^th^ percentiles mark the bottom and top of the box, respectively. Whiskers extend to the limits of the 95% confidence intervals. Scattered dots represent individual data points. Dashed lines indicate the values of HG-Ⅰ. (b) Linear relationship between genome size and genomic repeat content. The dashed line represents the best-fitting linear regression model. The grey zone indicates the 95% confidence intervals for the regression line. The adjusted *R*^2^ and *P*-value quantify the strength and significance of the relationship, respectively.

Plant of HG-Ⅰ owned a genome size as 840.082 Mb, deviated from which samples of the northern haplogroups (HG-Ⅱ, −Ⅳ, and −Ⅴ) had significantly smaller genome sizes (mean ±SD = 802.672 ±18.250 Mb; one-sample Wilcoxon test, *V* = 13, *P* = 5.002e-12) (Figure 3a). In contrary, plants of the southern haplogroups (i.e., HG-Ⅲ) owned relatively most variable genome sizes (mean ±SD = 837.624 ±31.595 Mb), nearly equal to that of HG-Ⅰ (one-sample Wilcoxon test, *V* = 82, *P* = 0.899) and significantly higher than those of the northern haplogroups (Wilcoxon rank sum exact test, *W* = 121, *P* = 1.222e-05). In summary, plants of HG-Ⅲ located in relatively lower latitudes including areas across the Tropic of Cancer (23.436°N, covering 55% of HG-Ⅲ accessions; Figure 1a), are prone to have larger genome sizes than that of the ancestral and other derived haplogroups in *R. indica* (Figures 2‒3).

### Ecological and genomic drivers of genome size variation

Phylogeny-corrected analysis revealed correlation between genome size variation and extrinsic environmental or intrinsic genomic variables. Populations with larger genomes were significantly associated with higher winter temperature (bio11) or lower temperature seasonality (bio4) characterized a tropical environment (Pearson’s coefficient *r* = 0.61 or −0.58, *P* = 1.200e-09 or 1.800e-08; PGLS adjusted *R*^2^ = 0.264 or 0.276, *P* = 0.035 or 0.031) (Table 1). Further investigation revealed a significant correlation between genome size variation and the content of specific genomic repeats (Figure 3b), particularly the 45S rDNA and *Ty1*-*copia* type long tandem repeats (PGLS adjusted *R*^2^ = 0.155 or 0.261, *P* = 0.039 or 0.036) (Table 2).

**Table 1.** Statistics for PGLS models detecting the significant effects of bioclimatic variables on genome size variation across populations of *R. indica*.

| Hierarchical group | df | AIC | *F*-statistic | Adjusted $R^2$ | *P*-value |
| --- | --- | --- | --- | --- | --- |
| Temperature seasonality/ bio4 | 2 | -73.177 | 5.953 | 0.276 | **0.031** |
| Mean temperature of coldest quarter/ bio11 | 2 | -72.939 | 5.650 | 0.264 | **0.035** |
| Isothermality/ bio3 | 2 | -72.803 | 5.480 | 0.256 | **0.037** |
| Minimum temperature of coldest month/ bio6 | 2 | -72.796 | 5.471 | 0.256 | **0.037** |
| Annual mean temperature/ bio1 | 2 | -72.680 | 5.327 | 0.250 | **0.040** |

**Table 2.** Statistics for PGLS models detecting the significant effects of genomic repeat variables on genome size variation across populations of *R. indica*.

| Hierarchical group | df | AIC | F-statistic | Adjusted $R^2$ | P-value |
| --- | --- | --- | --- | --- | --- |
| rDNA |  |  |  |  |  |
| 45S | 2 | -71.016 | 3.384 | 0.155 | **0.039** |
| 5S | 2 | -68.053 | 0.398 | 0.049 | 0.540 |
| Class I/ *Ty1-copia* |  |  |  |  |  |
| *Bianca* | 2 | -72.887 | 5.585 | 0.261 | **0.036** |
| *SIRE* | 2 | -69.811 | 1.183 | 0.014 | 0.298 |
| *TAR* | 2 | -67.918 | 0.330 | 0.054 | 0.576 |
| *Ivana* | 2 | -67.741 | 0.176 | 0.068 | 0.683 |
| *Tork* | 2 | -67.722 | 0.159 | 0.069 | 0.697 |
| *Ale* | 2 | -67.652 | 5.133 | 0.069 | 0.732 |
| Class I/ *Ty3-gypsy* |  |  |  |  |  |
| *Tekay* | 2 | -70.489 | 2.816 | 0.123 | 0.119 |
| *Retand* | 2 | -68.384 | 0.776 | 0.018 | 0.396 |
| *CRM* | 2 | -67.577 | 0.034 | 0.080 | 0.858 |
| *Athila* | 2 | -67.576 | 0.033 | 0.080 | 0.859 |
| *Galadriel* | 2 | -67.546 | 0.008 | 0.083 | 0.932 |
| Class I/ *LINE* | 2 | -67.608 | 0.060 | 0.078 | 0.810 |
| Class I/ Total | 2 | -67.597 | 0.051 | 0.079 | 0.825 |
| Class II/ *TIR* |  |  |  |  |  |
| *hAT* | 2 | -68.918 | 1.243 | 0.018 | 0.287 |
| *MuDR Mutator* | 2 | -68.502 | 0.855 | 0.011 | 0.373 |
| *PIF Harbinger* | 2 | -67.684 | 0.126 | 0.072 | 0.728 |
| *EnSpm CACTA* | 2 | -67.538 | 0.000 | 0.083 | 0.986 |
| Others/ *Satellite* | 2 | -68.971 | 1.294 | 0.022 | 0.278 |

### Fitness effect of genome size variation

The common garden experiment provided evidence for local adaptation of genotypes with varied genome sizes. Significant environment/site effects were detected in several traits, such as seed germination rate, survival or growth rate of seedlings, and plant architectures (Table 3; Figure S5). Under the temperate Kunming (KM), both seed germination rates and seedling survival rates were higher than those in the tropical Xishuangbanna (BN) (Wilcoxon rank sum exact tests, *P* ≤ 0.0001). Effect of genome size/GS type was only observed on days of flowering (Chisq = 11.807, *P* = 5.900e-04), with plants of larger genomes flowering much earlier than those of smaller genomes in either BN (mean ±SD of large vs. small, 90.572 ± 25.773 vs. 109.941 ±28.762; *P* = 0.002) or KM (212.826 ±43.690 vs. 234.917 ± 28.389; *P* = 0.001) (Figure S5). Significant two-way interactions of site ×GS on trait variation were observed in several fitness-related components, such as the relative growth rate and total number of fruits, under the relatively stable tropical environment of BN (Chisq = 11.587 and 9.400, *P* = 1.139e-03 and 2.170e-03) (Table 3). For example, accessions with larger genomes displayed improved performance in terms of lower relative growth rate and higher fecundity than those of accessions with small genomes in BN (Figure 4). The fitness advantage of larger genomes was clearly shown in BN rather than that in KM, exhibiting a pattern of conditional neutrality of genome size variation in southern populations/haplogroups of *R. indica*. In summary, larger genomes may confer an adaptive advantage under tropical conditions.

**Table 3.**
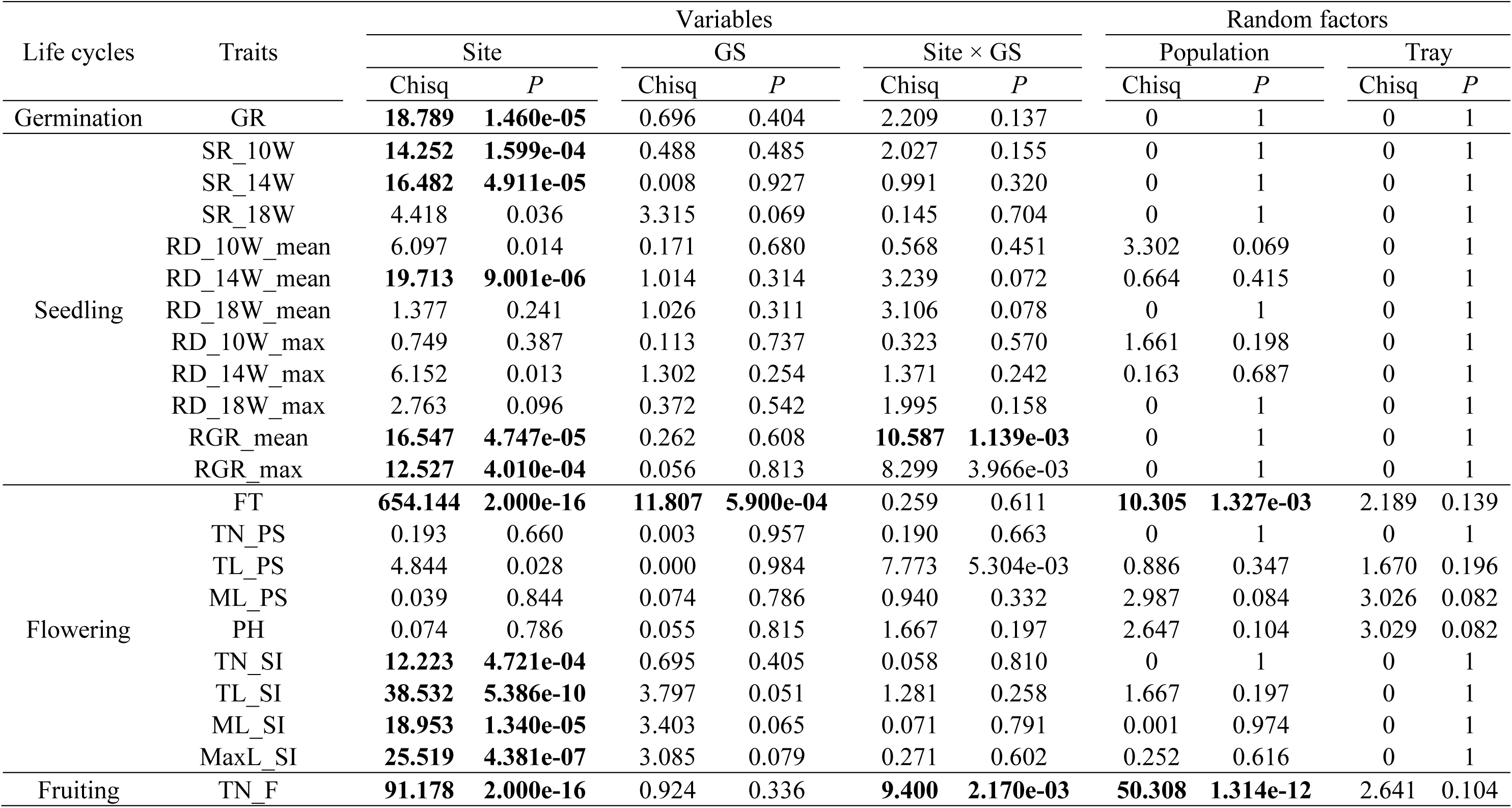
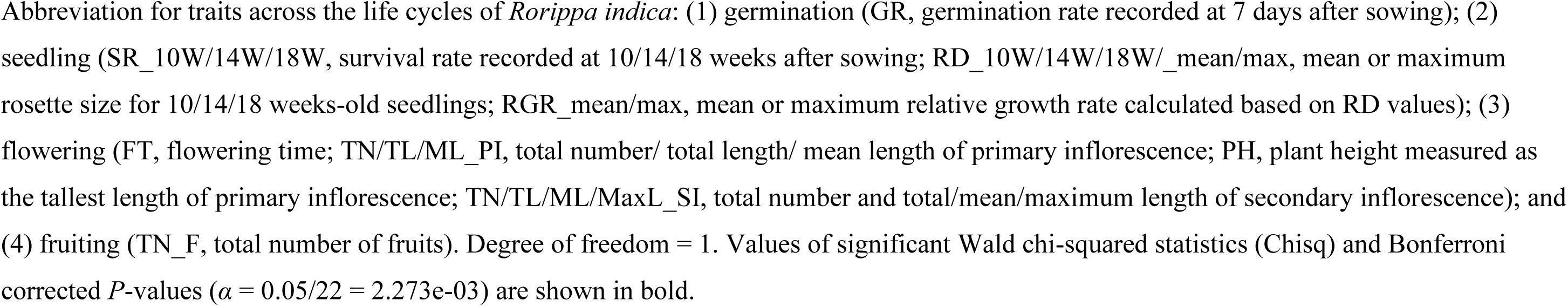
Summary of the GLMM analysis for detecting local adaptation of genome size variation in *R. indica*.

| Life cycles | Traits | Variables |  |  |  |  |  | Random factors |  |  |  |
| --- | --- | --- | --- | --- | --- | --- | --- | --- | --- | --- | --- |
| | | Site | | GS | | Site × GS | | Population | | Tray | |
|  |  | Chisq | *P* | Chisq | *P* | Chisq | *P* | Chisq | *P* | Chisq | *P* |
| Germination | GR | **18.789** | **1.460e-05** | 0.696 | 0.404 | 2.209 | 0.137 | 0 | 1 | 0 | 1 |
| Seedling | SR_10W | **14.252** | **1.599e-04** | 0.488 | 0.485 | 2.027 | 0.155 | 0 | 1 | 0 | 1 |
|  | SR_14W | **16.482** | **4.911e-05** | 0.008 | 0.927 | 0.991 | 0.320 | 0 | 1 | 0 | 1 |
|  | SR_18W | 4.418 | 0.036 | 3.315 | 0.069 | 0.145 | 0.704 | 0 | 1 | 0 | 1 |
|  | RD_10W_mean | 6.097 | 0.014 | 0.171 | 0.680 | 0.568 | 0.451 | 3.302 | 0.069 | 0 | 1 |
|  | RD_14W_mean | **19.713** | **9.001e-06** | 1.014 | 0.314 | 3.239 | 0.072 | 0.664 | 0.415 | 0 | 1 |
|  | RD_18W_mean | 1.377 | 0.241 | 1.026 | 0.311 | 3.106 | 0.078 | 0 | 1 | 0 | 1 |
|  | RD_10W_max | 0.749 | 0.387 | 0.113 | 0.737 | 0.323 | 0.570 | 1.661 | 0.198 | 0 | 1 |
|  | RD_14W_max | 6.152 | 0.013 | 1.302 | 0.254 | 1.371 | 0.242 | 0.163 | 0.687 | 0 | 1 |
|  | RD_18W_max | 2.763 | 0.096 | 0.372 | 0.542 | 1.995 | 0.158 | 0 | 1 | 0 | 1 |
|  | RGR_mean | **16.547** | **4.747e-05** | 0.262 | 0.608 | **10.587** | **1.139e-03** | 0 | 1 | 0 | 1 |
|  | RGR_max | **12.527** | **4.010e-04** | 0.056 | 0.813 | 8.299 | 3.966e-03 | 0 | 1 | 0 | 1 |
| Flowering | FT | **654.144** | **2.000e-16** | **11.807** | **5.900e-04** | 0.259 | 0.611 | **10.305** | **1.327e-03** | 2.189 | 0.139 |
|  | TN_PS | 0.193 | 0.660 | 0.003 | 0.957 | 0.190 | 0.663 | 0 | 1 | 0 | 1 |
|  | TL_PS | 4.844 | 0.028 | 0.000 | 0.984 | 7.773 | 5.304e-03 | 0.886 | 0.347 | 1.670 | 0.196 |
|  | ML_PS | 0.039 | 0.844 | 0.074 | 0.786 | 0.940 | 0.332 | 2.987 | 0.084 | 3.026 | 0.082 |
|  | PH | 0.074 | 0.786 | 0.055 | 0.815 | 1.667 | 0.197 | 2.647 | 0.104 | 3.029 | 0.082 |
|  | TN_SI | **12.223** | **4.721e-04** | 0.695 | 0.405 | 0.058 | 0.810 | 0 | 1 | 0 | 1 |
|  | TL_SI | **38.532** | **5.386e-10** | 3.797 | 0.051 | 1.281 | 0.258 | 1.667 | 0.197 | 0 | 1 |
|  | ML_SI | **18.953** | **1.340e-05** | 3.403 | 0.065 | 0.071 | 0.791 | 0.001 | 0.974 | 0 | 1 |
|  | MaxL_SI | **25.519** | **4.381e-07** | 3.085 | 0.079 | 0.271 | 0.602 | 0.252 | 0.616 | 0 | 1 |
| Fruiting | TN_F | **91.178** | **2.000e-16** | 0.924 | 0.336 | **9.400** | **2.170e-03** | **50.308** | **1.314e-12** | 2.641 | 0.104 |

Abbreviation for traits across the life cycles of *Rorippa indica*: (1) germination (GR, germination rate recorded at 7 days after sowing); (2) seedling (SR\_10W/14W/18W, survival rate recorded at 10/14/18 weeks after sowing; RD\_10W/14W/18W/\_mean/max, mean or maximum rosette size for 10/14/18 weeks-old seedlings; RGR\_mean/max, mean or maximum relative growth rate calculated based on RD values); (3) flowering (FT, flowering time; TN/TL/ML\_PI, total number/ total length/ mean length of primary inflorescence; PH, plant height measured as the tallest length of primary inflorescence; TN/TL/ML/MaxL\_SI, total number and total/mean/maximum length of secondary inflorescence); and (4) fruiting (TN\_F, total number of fruits). Degree of freedom = 1. Values of significant Wald chi-squared statistics (Chisq) and Bonferroni corrected *P*-values ( $\alpha = 0.05/22 = 2.273e-03$ ) are shown in bold.

**Fig. 4.**
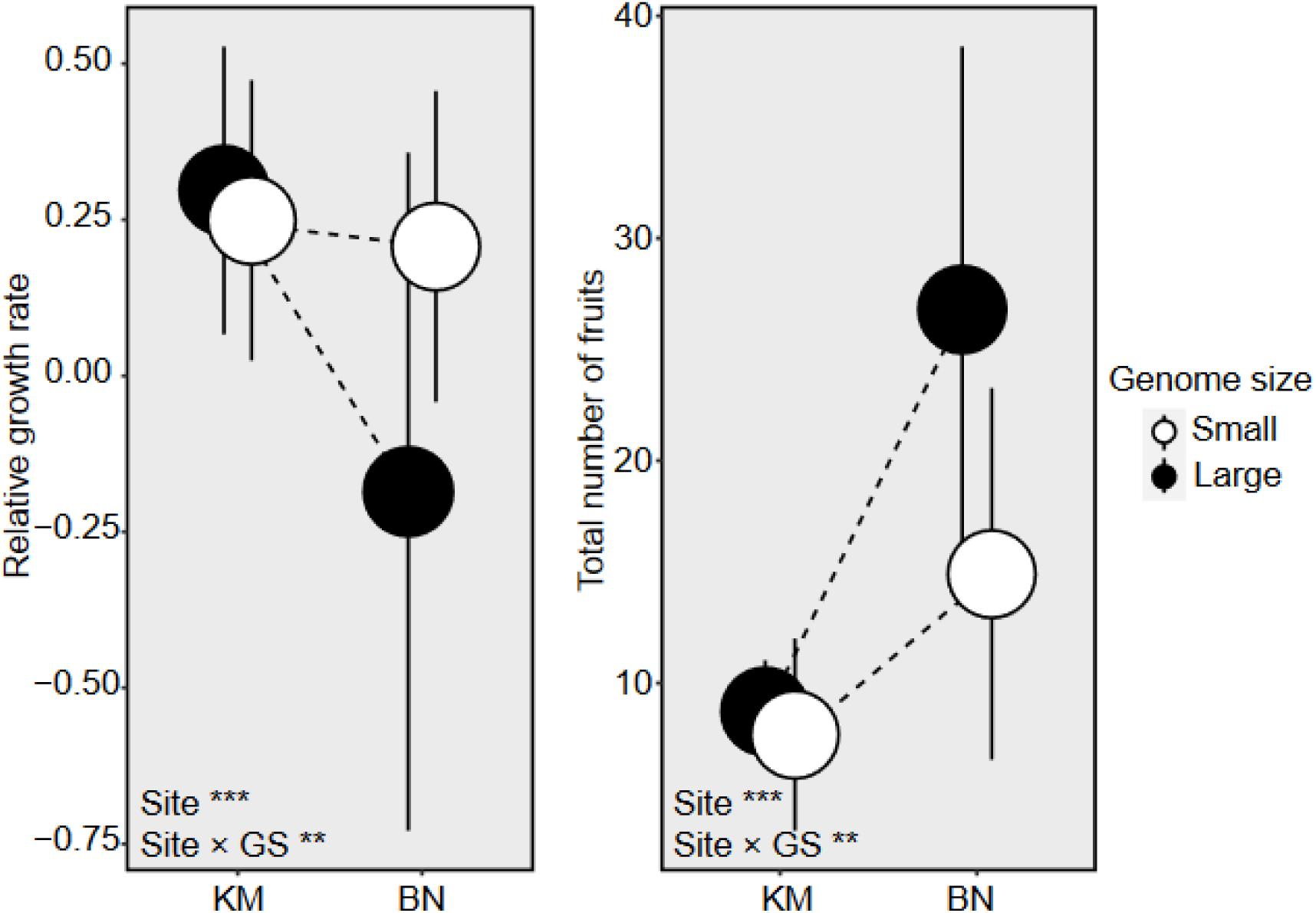
Local adaptation of larger genomes to tropical environment. Reaction norms of phenotypic variation in relative growth rate (left) and total number of fruits for small (open dots) and large genomes (closed dots) in two contrasting common gardens: Kunming (KM) and Xishuangbanna (BN). The dashed lines connect the mean values for each trait and genome size type across the two common gardens. Error bars represent the standard deviations. Asterisks indicate statistically significant differences between sites or interactions between site and genome size (site ×GS) (\*\**P* < 0.01, \*\*\**P* < 0.001), as determined by Bonferroni-corrected tests.

## Discussion

This study tests the “large genome selection” hypothesis by investigating the intraspecific genome size variation in a widespread plant *R. indica*. Our findings suggest that plant genome size may tend to be larger due to the expansion of repeat elements, particularly 45S rDNA and *Ty1*-*copia* type long tandem repeats. Such genomic enlargement is a conditionally adaptive response to seasonally temperature-stable environments, allowing for the accumulation of repetitive genetic material with potential fitness effect (Figure 5). The observed local adaptation in the reciprocal transplantation experiment further strengthens this argument. Overall, this study provides valuable insights into drivers and consequences of intraspecific genome size variation in *R. indica*, improving our understanding of genome evolution and adaptation in plants.

**Fig. 5.**
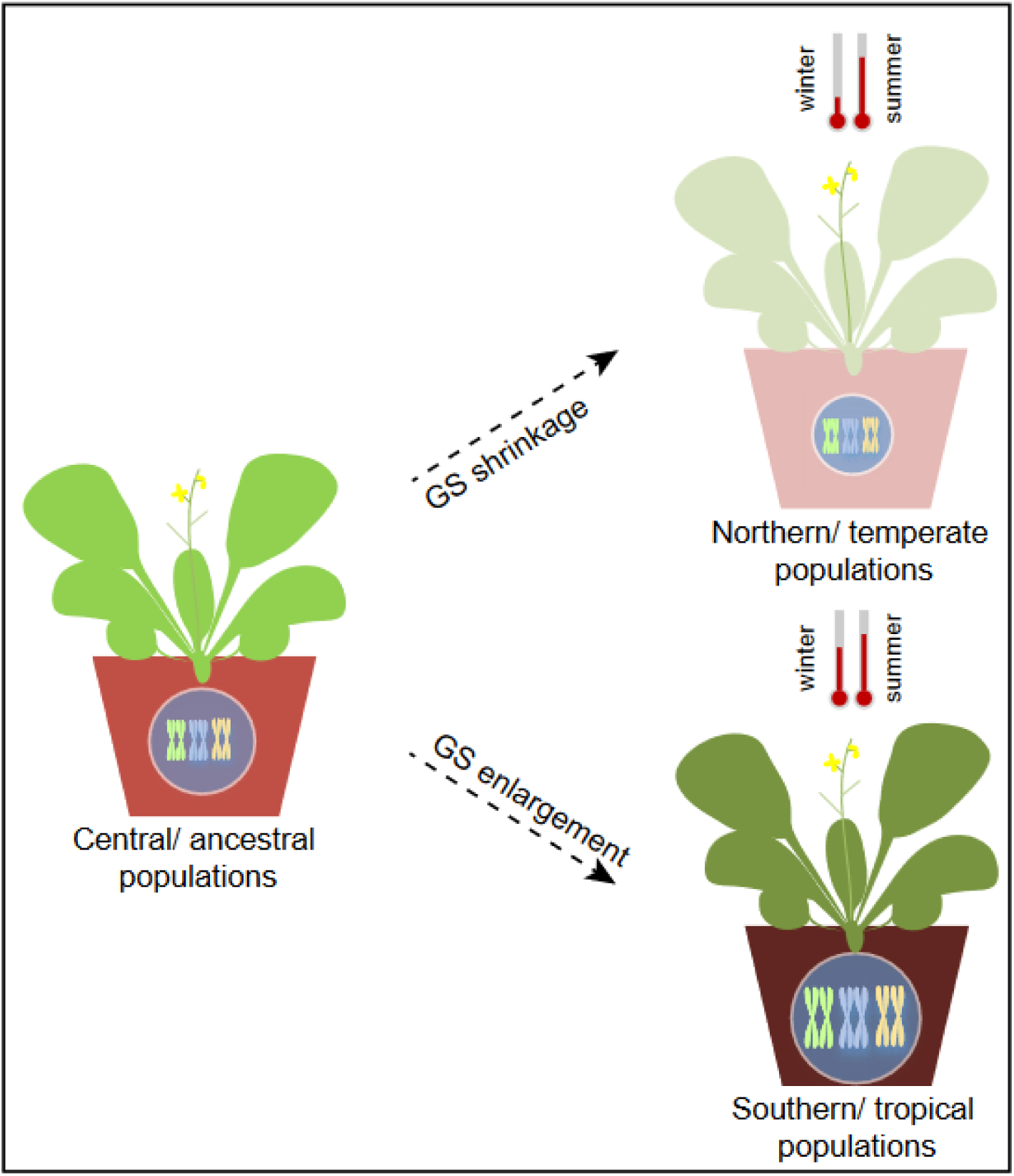
Evolutionary model of genome size variation in *R. indica*. The schematic illustration depicts the hypothesized evolutionary history of genome size variation of *R. indica* plants. It suggests that the species originated in central or mid-latitude regions with a moderate genome size (left). As the population expanded, natural variation in genome size (GS) emerged (right). Smaller genomes became more common in northern/temperate regions with higher temperature seasonality, while larger genomes predominated in southern/tropical regions with more stable temperatures. The nucleus represents the genome, with its size proportional to the chromosome/genome size. The thermometers above plants illustrate the seasonal temperature variation in each region.

### Natural variation in genome size of *R. indica*

We reported a substantial 128 Mb intraspecific variation among populations of *R. indica* (Figure 1), exceeding that of *Arabidopsis thaliana* (18‒23 Mb) (Long *et al*., 2013; Lian *et al*., 2024) and other relative *Rorippa* species (Han *et al*., 2022). The wide range of genome sizes suggested a high degree of cytological diversity in *R. indica*, which can be attributed to several evolutionary processes during the polyploid origin and establishment of *R. indica*. First, the allohexaploid origin made *R. indica* reshuffle its genome that incorporated parental species with different genome sizes. The potential maternal progenitor *R. dubia* is an allotetraploidy who also exhibits plentiful natural variation in genome size (Tu *et al*., 2019; Zheng *et al*., 2021; Han *et al*., 2024). This provides *R. indica* plants with opportunity to alter genome sizes from a hybrid and relatively larger genome founder, such as HG-Ⅰ (Figure 2). Second, polyploidy can trigger the so-called ‘genome shock’ and associated genome size changes through relaxed selection on duplicated genomic elements, including those repetitive ones, during DNA replication processes (McClintock, 1984; Cheng *et al*., 2018). Coincidence with the findings in *A. thaliana* and maize studies (Long *et al*., 2013; Bilinski *et al*., 2018), larger genomes in *R. indica* are associated with an expansion of repeat elements, particularly 45S rDNA and *Ty1*-*copia* type long tandem repeats, after accounting for the influence of phylogeny (Table 2). This suggests repetitive or transposable elements play crucial roles in shaping genetic or phenotypic diversity within *R indica* (Gu & Hsu, 1986; Tu *et al*., 2019). Third, plants of *R. indica* established clear phylogeographic patterns very recently around one million years ago (Figures 2‒3). These quickly differentiated populations may incur divergent resolution through independent gain or loss of genomic elements (Taylor *et al*., 2001), permitting plentiful intraspecific genome size variation to be generated and maintained in face of limited polymorphism (Lefebure *et al*., 2017). Intriguingly, genome size variation in *R. indica* is linked to geographic location and temperature (Figure 1b; Table 1), indicating a potential adaptive role in different environments.

### Temperature as a significant driver of genome size variation in *R. indica*

Intraspecific genome size variation is a widespread phenomenon observed across many taxa (Šmarda & Bureš, 2010; Díez *et al*., 2013; Bilinski *et al*., 2018). Understanding the factors driving this variation is crucial for comprehending the evolutionary dynamics of species and their adaptation to diverse environmental conditions. This study investigates the potential relationship between intraspecific genome size variation and environmental variables in *R. indica*. Our analysis revealed that environmental variables, particularly temperature, are significant determinants of genome size variation in *R. indica* (Figure 1b; Table 1). These findings suggest that genome size variation can play a role in adaptation to different environments experienced by plants within this species (Faizullah *et al*., 2021).

Previous studies revealed a complex relationship between genome size variation and temperature variables. In some cases, species found in colder, higher-latitude regions tend to have larger genomes, such as the Antarctic krill (Shao *et al*., 2023) or marine microbes (Ngugi *et al*., 2023). This may be attributed to the slower metabolic rates and cellular processes required for survival in colder environments. However, this pattern is not universal and can vary across organisms (e.g., plants) with different life history strategies (Simonin & Roddy, 2018; Bureš *et al*., 2024). In contrast, larger genomes may be advantageous in warmer, wetter and more stable environments, such as the cases in *Helianthus* sunflowers (Qiu *et al*., 2019), wild maize (Díez *et al*., 2013), tropical gingers (Zingiberaceae) (Xavier *et al*., 2024), and lily family (Liliaceae) (Carta & Peruzzi, 2016). As stated in the “large genome selection” hypothesis, this could be attributed to large-genome mediated factors like increased genetic diversity, enhanced stress tolerance, or altered developmental patterns, upon which novel raw genetic materials can be provided for natural selection to act (Bennett, 1987; Lefebure *et al*., 2017). For example, larger genomes may be advantageous due to relaxed selection pressures on stomatal size or growth rates, enabling the accumulation of biomass and facilitating fecundity (Schley *et al*., 2022). Furthermore, in warmer or more stable environments, the activity of transposable element would be spurred and hard to be purged (Kelly *et al*., 2015), relaxing “large genome constraint” and promoting new genetic or phenotypic variation (Kelly *et al*., 2015; Lefebure *et al*., 2017). Specific repeat elements were also identified as significant drivers of genome size variation and contributed to stressful adaptation (Schley *et al*., 2022). These elements can potentially influence the adaptive performance of species in different thermal environments. While our findings provide compelling evidence for a relationship between genome size and temperature in *R. indica*, it is important to acknowledge that the relationship is complex and influenced by multiple factors (Díez *et al*., 2013). Incongruence or mixed relationship also existed between intra- and interspecific studies (e.g., Cacho *et al*., 2021), suggesting the necessity of accounting for phylogenetic autocorrelation or population demographic interference (Table 1; Figures 2‒3), as well as the range of environmental gradients for focal systems. Furthermore, experimental studies are needed to elucidate the underlying fitness effects linking genome size variation to temperature-related adaptations (Table 3; Figure 4).

### Fitness effects of genome size variation

Our study underscores the potential fitness consequences of genome size variation in *R. indica*. We found that accessions with larger genomes exhibited superior performance in certain fitness traits like the total number of fruits, conditionally under tropical environments (Figure 4). This suggests that sizable genomes may provide fitness advantages that enable organisms to thrive in diverse challenging conditions, such as the newly colonized tropical areas by *R. indica* (Setyawati *et al*., 2015). First, the observed fitness benefits of larger genomes in tropical environments may be linked to their increased genetic diversity or genomic plasticity. Larger genomes often contain more repetitive elements, which can provide a reservoir of genetic material for recombination and mutation (Mei *et al*., 2018; Schley *et al*., 2022). This increased genetic variation can further facilitate adaptation to changing environmental conditions, such as higher temperature, precipitation, or herbivory. Second, larger genomes may possess more complex regulatory networks, allowing for finer control of gene expression or epigenetic modification. This could be particularly advantageous in tropical environments, where organisms must respond to rapidly changing conditions or cope with a wide range of biotic and abiotic stressors (Satake *et al*., 2022). For example, larger genomes might contain more regulatory genes or microRNAs that can modulate gene expression in response to environmental cues (Leitch & Leitch, 2008). Third, those so-called constraints on larger genomes would be relaxed given the relatively unrestricted energy or resource supply in the tropics. While larger genomes offer benefits, they may also come with potential costs, for example, requiring more energy to replicate or exhibiting more susceptibility to deleterious mutations (Petrov, 2002; Lynch & Conery, 2003; Francis *et al*., 2008). The optimal genome size may therefore depend on the specific environmental challenges faced by southward *R. indica* lineages and the trade-offs between the benefits and costs of genome size variation. Although the fitness effect of the larger genome was clearly unraveled, it may not fully capture the entire adaptive landscape of genome size variation within *R. indica*. Future studies could investigate the mechanisms and evolutionary context of genome size variation by comparing *R. indica* with other species.

### Conclusion

By investigating the natural variation of genome size in *R. indica*, we have provided insights into the potential drivers of this enigmatic phenomenon. Our study highlights the role of repeat element expansion and suggests a link between genome size increase and adaptation to temperature-stable environments in the Tropics, thus supporting the “large genome selection” hypothesis. This research paves the way for further exploration of the interplay between genome size and organismal fitness in the face of environmental pressures.

## Author Contributions

T-SH and Y-WX conceived the project. T-SH collected the data, performed analyses and drafted the manuscript. Q-JZ performed the flow cytometry experiment and phylogenetic network analysis. J-XL and SL participated in the common garden experiments. Z-QD analyzed the phenotypic data. All authors reviewed the final manuscript.

## Acknowledgements

This work was supported by the National Natural Science Foundation of China (32170224 and 32225005), the NSFC-ERC International Cooperation and Exchange Programs (32311530331), and the Youth Innovation Promotion Association CAS (2020391). We acknowledge Prof. Hang Wang at the Southwest Forestry University for assisting common garden experiment.

## Data Availability

The sequences can be accessed by GenBank IDs: PQ433226‒PQ433289.

## Supporting Information

The following Supporting Information is available for this article:

**Figure S1** FCM-estimated genome size variation in *R. indica* and its relatives.

**Figure S2** Correlation between genome size and bioclimatic or geographic variables.

**Figure S3** Bayesian tree of chloroplast haplotypes in *R. indica*.

**Figure S4** Experimental test of the fitness effect of larger genomes.

**Figure S5** Traits collected for plants with different genome sizes.

**Table S1** PGLS statistics for the ecological effects on genome size.

## Notes

### Competing Interest Statement

The authors have declared no competing interest.

### Summary of Updates

In the revised manuscript, we have made the Latin numerals more prominent by providing a PDF version with enhanced formatting. Additionally, we have clarified the institutional affiliations for all authors. Figure 1 has also been refined to enhance its clarity. No other changes have been made to the content.

